## Supporting Information for "Simulated hemiparesis increases optimal spatiotemporal gait asymmetry but not metabolic cost"

### Introduction

OpenSim Moco contains several built-in goals which, when added to an optimization, act as objective functions or constraints. The goal to minimize the sum of integrated muscle excitations cubed was already implemented within the Moco software, but we had to develop the goals to minimize step length and step time asymmetry for our project. The following sections summarize the development of step length and step time symmetry goals that we built for this study, which are both now implemented in the release of Moco included with OpenSim 4.3.

### Step Length Symmetry Goal:

This goal was developed to be added to a Moco optimization to minimize the difference between the step length asymmetry of the gait model and a user-specified target asymmetry value. The step length asymmetry (SLA) is computed as a ratio of the right step length ($SL_R$) minus the left step length ($SL_L$) divided by the $SL_R$ plus the $SL_L$ (Eq. A1), note that for our project the right limb is the non-paretic side and the left limb is the paretic side.

$$SLA = \frac{SL_R - SL_L}{SL_R + SL_L} \times 100 \qquad \text{(A1)}$$

The right step length is defined as the anterior-posterior distance between the right and left heel contact elements at the instant of right foot-strike (defined based on a threshold level of ground reaction force). Ideally, we would want to be able to compute the step length for each limb for the current iteration of the optimization and use those values to compute the step length asymmetry. However, the *MocoGoal* interface only allows computing cost values integrated across the trajectory or values at the endpoint of the trajectory, and values relevant to asymmetry (i.e., heel strikes) can occur at intermediate points in the trajectory. Therefore, we needed to develop a way to approximate the step length asymmetry without using the actual step lengths, since these would only be computed at discrete points in time (at the instant of right and left heel strike).

Therefore, in this goal, called *MocoStepLengthAsymmetryGoal*, we use the distances between the heel contact elements to approximate the step length for each limb. To do this, the user first inputs a total stride length for the gait cycle and the target SLA (TargetSLA) value. A symmetric step length is set when the target SLA value is equal to zero. The goal then computes the right step length and left step length, and places bounds on the distance between right (RightStepBound) and left (LeftStepBound) heel contact elements (Eq. A2) to add a penalty to the objective function that accumulate when the distance between these heel elements exceeds the bounds.

$$RightStepBound = \frac{1}{2}(1 - TargetSLA) * StrideLength$$
$$LeftStepBound = \frac{1}{2}(1 + TargetSLA) * StrideLength \tag{A2}$$

Once the optimization is ongoing, for each iteration and at each time node (t), the goal computes the distance between the right (RightHeelPos) and left (LeftHeelPos) heel contact element positions at time t, and compares them to the bounds we set for the goal.

$$RightDistance(t) = RightHeelPos(t) - LeftHeelPos(t) \tag{A3}$$
$$LeftDistance(t) = LeftHeelPos(t) - RightHeelPos(t)$$

We use a smoothing function to help make the function smooth across all possible distances between heels, with a smoothing ($S_1$) term that dictates the shape of the smoothing function (Eq. A4).

$$\mathrm{RightFootPenalty(t)} = 0.5 + \frac{1}{2} * -\tanh\big(\mathrm{S_1} * (\mathrm{RightStepBound} - \mathrm{RightDistance(t)})\big)$$
$$\mathrm{LeftFootPenalty(t)} = 0.5 + \frac{1}{2} * -\tanh(\mathrm{S_1} * (\mathrm{LeftStepBound} - \mathrm{LeftDistance(t)})) \tag{A4}$$

Finally, these penalty terms for the right and left side get added together to form the integrand. This process gets repeated for every time point throughout the motion and then these penalties are integrated over time to result in the objective function value for the step length goal. Since this is an approximation of SLA, we compute the actual SLA for the optimal solution and compare it against our target SLA. If these values deviate beyond an acceptable error, we can adjust settings (such as the goal weighting, the smoothing term, or the target SLA itself) to try to get an optimal result closer to our target asymmetry. For more information on the goal, users can refer to the OpenSim Moco source code.

**Step Time Symmetry Goal:**

As above, we developed this goal for implementation within the Moco software framework to minimize the difference between the model's step time asymmetry (STA) and a user-defined target asymmetry value, where a value of 0 would target a symmetrical step time. Step time is computed as the time between consecutive foot strikes, such that the right step time ($ST_R$) is the time from left foot strike to right foot strike, and left step time ($ST_L$) is the time from right foot strike to left foot strike. Then, the step time asymmetry is the ratio of the right step time ($ST_R$) minus the left step time ($ST_L$) divided by the $ST_R$ plus the $ST_L$ (Eq. A5).

$$STA = \frac{ST_R - ST_L}{ST_R + ST_L} \times 100 \tag{A5}$$

Just like for the step length goal, we cannot compute the step time asymmetry in the ideal way. Therefore, we developed an approximate form of step time asymmetry for use within direct collocation. For this goal, the step time asymmetry is computed by counting the number of nodes that each foot is in contact with the ground based on a threshold vertical ground reaction force value. Since walking contains two double support phases where both feet are on the ground, when the goal detects that both feet are on the ground, it must also detect which foot is in front and it then assigns the step time to the leading foot. Overall, this goal estimates the time between consecutive foot strikes to approximate the left and right step times.

First, users define the target STA value for the goal. Once the optimization is running, for each iteration and at each time node (t), the function detects if the right foot is in contact with the ground (RightContactDetect) and if the left foot is in contact with the ground (LeftContactDetect) using the ground reaction force (RightGRF(t) or LeftGRF(t)) and the foot strike threshold value (Threshold) with a smoothing parameter ($S_2$) (Eq. A6).

$$RightContactDetect(t) = -1 * \left(0.5 + \left(\frac{1}{2} * \tanh\left(S_2 * RightGRF(t) - Threshold\right)\right)\right) \quad \text{(A6)}$$

$$LeftContactDetect(t) = -1 * \left(0.5 + \left(\frac{1}{2} * \tanh\left(S_2 * LeftGRF(t) - Threshold\right)\right)\right)$$

The right foot is assigned a negative value such that shorter right step times give negative asymmetry values (as in Eq. A5). Then, the goal detects which of the feet are the most forward in the anterior-posterior direction, using the positions of the heel contact element on each foot. We use another smoothed function with a smoothing parameter ($S_3$) that is designed to be -1 when the left foot is in front and +1 when the right foot is in front (Eq. A7).

$$FrontFoot(t) = \tanh\left(S_3 * \left(RightHeelPos(t) - LeftHeelPos(t)\right)\right) \quad \text{(A7)}$$

Then, we use the value calculated in A7 with a 'Tie Breaker' function (Eq. A8) which is designed to be 0 when the model is in single support, +1 when the model is in double support with the left heel in front, and -1 when the model is in double support with the right heel in front.

$$TieBreaker(t) = RightContactDetect(t) * LeftContactDetect(t) * FrontFoot(t) \quad \text{(A8)}$$

The value of TieBreaker is then used to assign the current time node (t) to either a left step or a right step with an equation that corrects for the double support phase that we determined in eq. A8, where if the model is in the left step phase (Eq. A9).

$$\begin{aligned} LeftStep(t) &= LeftContactDetect(t) + TieBreaker(t) \\ RightStep(t) &= RightContactDetect(t) + TieBreaker(t) \end{aligned} \quad \text{(A9)}$$

Finally, we use another smooth function (Eq. A10) that becomes the integrand for the time point (t) with a smoothing term ($S_4$).

$$Integrand(t) = \tanh\left(S_4 * \left(LeftStep(t) + RightStep(t)\right)\right) \quad \text{(A10)}$$

The integrand value gets integrated over time and then compared against the target STA value that the user inputs. The error between the integrand value and the target STA value is squared and used in the weighted objective function.

As with the step length goal, this value approximates the actual step time asymmetry, therefore after the optimization terminates, we check the real value and adjust settings to the objective function weighting, smoothing terms, or foot contact threshold as needed to try to get a result with a STA close to our target value. For more information on the goal, users can refer to the OpenSim Moco source code.
